# Antimicrobial functions of lactoferrin promote genetic conflicts in ancient primates and modern humans

**DOI:** 10.1101/043091

**Authors:** Matthew F. Barber, Zev Kronenberg, Mark Yandell, Nels C. Elde

## Abstract

Lactoferrin is a multifunctional mammalian immunity protein that limits microbial growth through sequestration of nutrient iron. Additionally, lactoferrin possesses cationic protein domains that directly bind and inhibit diverse microbes. The implications for these dual functions on lactoferrin evolution and genetic conflicts with pathogens remain unclear. Here we show that lactoferrin has been subject to recurrent episodes of positive selection during primate divergence predominately at antimicrobial peptide surfaces consistent with long-term antagonism by pathogens. An abundant lactoferrin polymorphism in human populations and Neanderthals also exhibits signatures of positive selection across primates, linking ancient host-microbe conflicts to modern human genetic variation. Rapidly evolving sites in lactoferrin further correspond to molecular interfaces with pathogenic bacteria causing meningitis, pneumonia, and sepsis. Because microbes actively target lactoferrin to acquire iron, we propose that the emergence of antimicrobial activity provided a pivotal mechanism of adaptation sparking evolutionary conflicts via acquisition of new protein functions.

**AUTHOR SUMMARY:** Immunity genes can evolve rapidly in response to antagonism by microbial pathogens, but how the emergence of new protein functions impacts such evolutionary conflicts remains unclear. Here we have traced the evolutionary history of the lactoferrin gene in primates, which in addition to an ancient iron-binding function, acquired antimicrobial peptide activity in mammals. We show that, in contrast to the related gene transferrin, lactoferrin has rapidly evolved at protein domains that mediate iron-independent antimicrobial functions. We also find evidence of natural selection acting on lactoferrin in human populations, suggesting that lactoferrin genetic diversity has impacted the evolutionary success of both ancient primates and humans. Our work demonstrates how the emergence of new host immune protein functions can drastically alter evolutionary conflicts with microbes.

## INTRODUCTION

Genetic conflicts between microbial pathogens and their hosts are a major source of evolutionary innovation [1]. Selective forces imposed by these antagonistic interactions can give rise to dramatic bouts of adaptive gene evolution through positive selection. J.B.S. Haldane originally speculated on the importance of infectious disease as an “evolutionary agent” over 60 years ago [2], and the Red Queen hypothesis later posited that predators and their prey (or pathogens and their hosts) must constantly adapt in order to sustain comparative fitness [3,4]. More recent studies have demonstrated how evolutionary conflicts progress at the single gene or even single nucleotide level, as molecular interfaces between host and microbial proteins can strongly impact virulence and immunity [5–7]. Host-pathogen interactions thus provide fertile ground for studying rapid gene evolution and acquisition of novel molecular traits [8].

Lactoferrin presents a compelling model for investigating adaptation from an ancestral “housekeeping” function to a specialized immunity factor. Lactoferrin arose from a duplication of the transferrin gene in the ancestor of eutherian mammals roughly 160 million years ago [9]. A fundamental and shared feature of these proteins is the presence of two evolutionary and structurally homologous iron binding domains, the N and C lobes, each of which chelates a single iron ion with high affinity. Iron binding by these proteins can effectively starve invasive microbes of this crucial metal, a protective effect termed nutritional immunity [10,11]. Pathogens in turn actively scavenge iron from these and other host proteins in order to meet their nutrient requirements [12,13]. The importance of iron in human infectious disease is highlighted by genetic disorders of iron overload, such as hereditary hemochromatosis, which render affected individuals highly susceptible to bacterial and fungal infections [14,15]. In addition to its role in nutritional immunity, lactoferrin has acquired new immune functions independent of iron binding following its emergence in mammals. Lactoferrin is expressed in a variety of body tissues and fluids including breast milk, colostrum, saliva, tears, mucous, as well as the secondary granules of neutrophils and inhibits the growth of many microbes [16]. Portions of the lactoferrin N lobe are highly cationic, facilitating interaction with and disruption of microbial membranes. Two regions of the lactoferrin N lobe in particular, lactoferricin and lactoferrampin, can be liberated from the lactoferrin polypeptide by proteolytic cleavage and exhibit potent antimicrobial activity against bacteria, fungi, and viruses [17,18]. Lactoferrin, as well as lactoferricin alone, can directly bind the lipid A component of lipopolysaccharide (LPS) as well as lipoteichoic acid, contributing to interactions with surfaces of Gram-negative and Gram-positive bacteria [19,20]. Lactoferrin thus poses a unique challenge for pathogens – while its ability to bind iron makes it an attractive target for “iron piracy,” lactoferrin surface receptors could render these microbes more susceptible to associated antimicrobial activity. Despite a growing appreciation for lactoferrin’s immune properties, the evolutionary implications of these unique functions remain unclear. In the present study we decipher recent signatures of natural selection acting on lactoferrin in primates and modern humans to understand the evolutionary consequences of a newly acquired antimicrobial activity from a distinct ancestral function.

## RESULTS

### Positive selection has shaped the lactoferrin N lobe in primates

To assess the evolutionary history of lactoferrin in primates, we assembled gene orthologs from publicly available databases and cloned lactoferrin complementary DNA (cDNA) prepared from primary cell lines. In total, we compared 15 lactoferrin orthologs from hominoids, Old World, and New World monkeys, representing roughly 40 million years of primate divergence (Figure 1A, S1). We then used maximum likelihood-based phylogenetic approaches (performed with the PAML and HyPhy software packages) to calculate nonsynonymous to synonymous substation rate ratios (*d*N/*d*S) across this gene phylogeny [21–23]. Complementary tests indicated that lactoferrin has evolved under episodic positive selection in the primate lineage, consistent with a history of host-pathogen evolutionary conflict (Figure 1A, Supplemental Tables 1–7).

**Figure 1.**
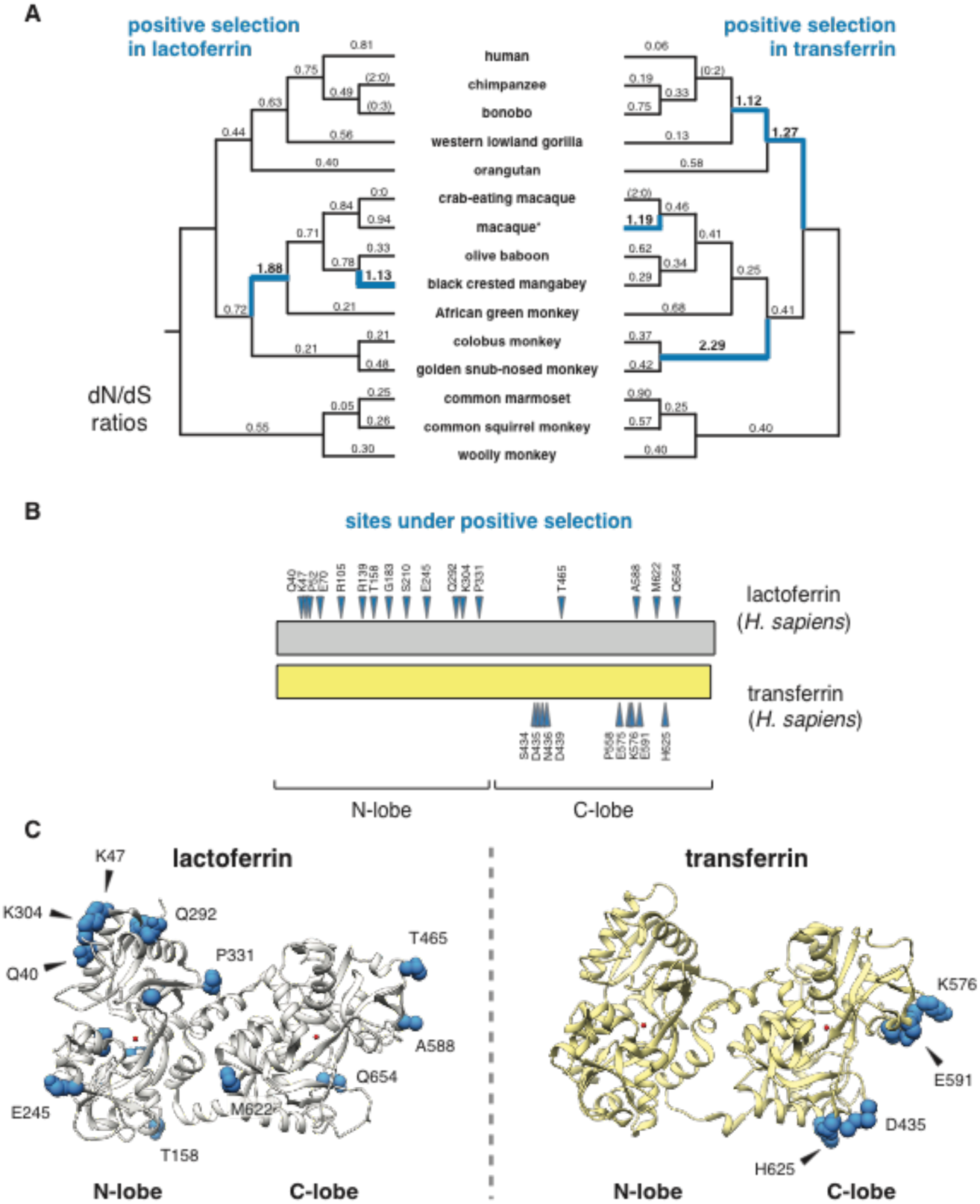
Dynamic evolution of the lactoferrin N lobe in primates. **A.** Paired primate phylograms showing signatures of positive selection in lactoferrin and transferrin. **d*N/*d*S* ratios along each lineage are shown, with ratios greater than 1 (indicative of positive selection) shown in blue. Branches with no silent or nonsynonymous mutations display ratios in parentheses. *For lactoferrin analyses the sequence of the Taiwanese macaque was used, whereas for transferrin rhesus macaque was included. This difference does not change the topology of the primate phylogram. **B.** Sites subject to positive selection in lactoferrin and transferrin are shown (blue arrows) along a schematic of the two proteins (phylogenetic analysis by maximum likelihood, posterior probability >0.95 by Naïve and Bayes Empirical Bayes analyses). The relative positions of the N and C lobes are shown. **C.** Ribbon diagrams for crystal structures of diferric lactoferrin (PDB: 1LFG) and transferrin (PDB: 3V83), with side chains of sites under positive selection calculated in **B** shown in blue. Iron in the N and C lobes is shown in red.

These findings are also in line with previous genome-wide scans for positive selection in primates which identified the lactoferrin gene (*LTF*) among others [24]. We next determined signatures of selection across individual codons in lactoferrin. In total, 17 sites displayed strong evidence of positive selection (posterior probability >0.95 from Naïve Empirical Bayes and Bayes Empirical Bayes analyses in PAML), with 13 of the 17 sites found in the N lobe (Figures 1B, 1C,S1,Supplementary Tables 2, 4–6). This observation was notably dissimilar from a parallel analysis of serum transferrin, where sites under positive selection were restricted to the C lobe (Figure 1B, 1C, Supplementary Table 3). These results are further consistent with our previous work indicating that rapid evolution in primate transferrin is likely due to antagonism by the bacterial iron acquisition receptor TbpA, which exclusively binds the transferrin C lobe [25–28]. Thus, while lactoferrin and transferrin both exhibit signatures of positive selection in primates, patterns of selection across the two proteins are highly discordant.

### Evolution and diversity of lactoferrin in modern humans

Evidence of episodic positive selection in primate lactoferrin led us to more closely investigate variation of this gene across human populations. Of the 17 sites we identified as rapidly evolving across primate species, amino acid position 47 overlapped with a high frequency arginine (R) to lysine (K) substitution in the N lobe of lactoferrin found in humans (Figure 2A, Supplementary Tables 8, 9). This position is markedly polymorphic between populations; while individuals of African ancestry carry the K47 allele at about 1% frequency, this variant is found in non-African populations at roughly 30-65% allele frequency, with the highest frequencies observed among Europeans (Figure 2B, Supplementary Table 9). The presence of R47 in related great apes combined with its high frequency in African populations suggests that R47 is the ancestral allele in humans, although it is conceivable that long-term balancing selection could also produce such differences. Data from the Neanderthal genome browser (http://neandertal.ensemblgenomes.org) further revealed lysine to be the consensus residue at position 47 in recently sequenced Neanderthals. The presence of the lactoferrin K47 allele in Neanderthal and non-African human populations and its near absence in Africans suggests one of several intriguing genetic models for the history of this variant, including long-term balancing selection, convergent evolution, or introgression from Neanderthals into modern humans.

**Figure 2.**
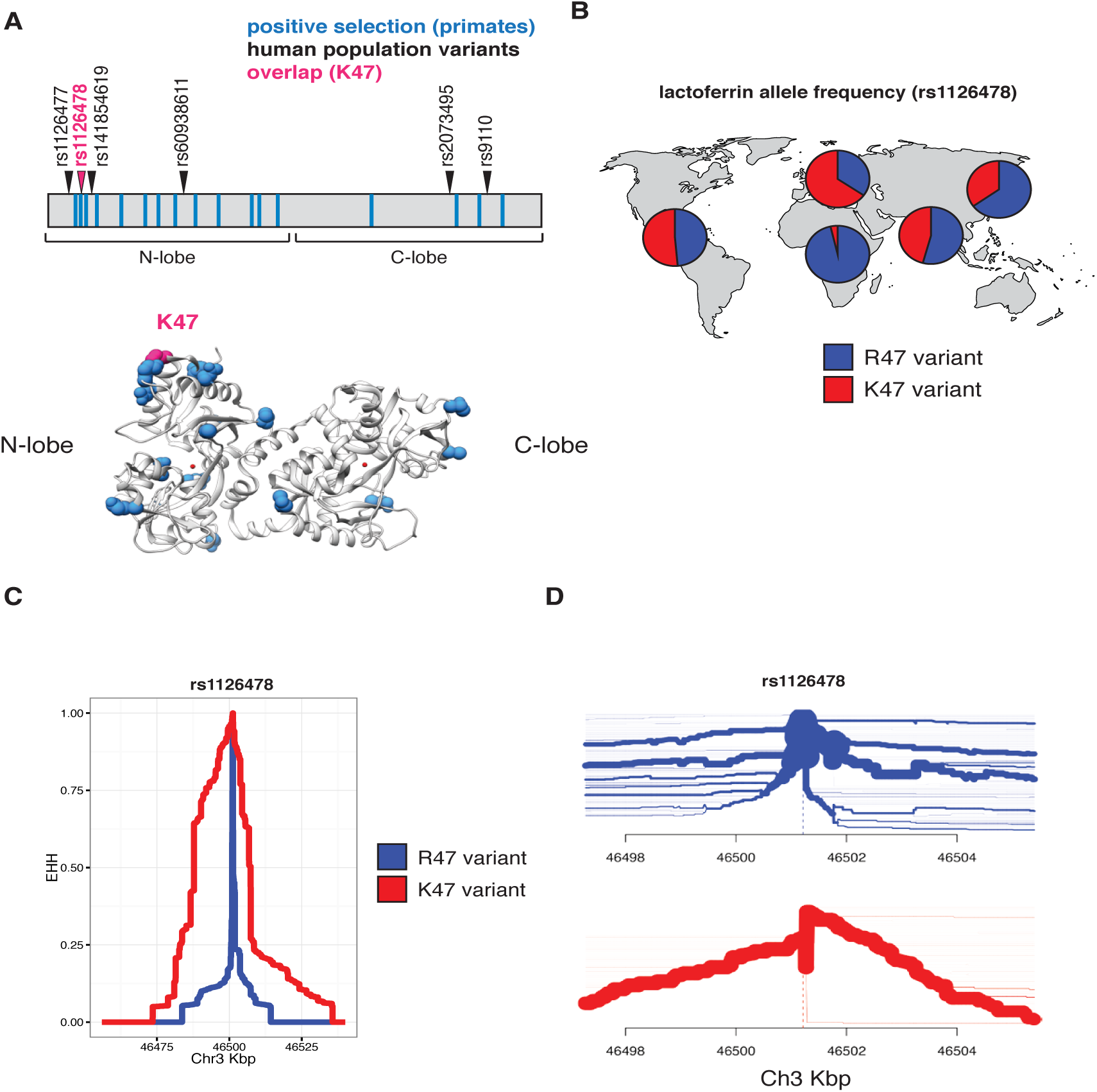
Diversity and evolution of human lactoferrin. **A.** Schematic representation of the lactoferrin protein showing positions of abundant (>1% allele frequency) nonsynonymous polymorphisms found in humans (arrows). Sites previously identified under positive selection across primates are shown as blue bars. The position of one variant, rs1126478 at amino acid position 47, which is also rapidly evolving in primates, is shown in magenta. The position of lysine 47 (K47) is also shown in the lactoferrin crystal structure (bottom panel). **B.** Relative allele frequencies of the R47 (blue) and K47 (red) lactoferrin variants shown as pie charts across human populations. Data were obtained from the 1000 Genomes Project Phase III. **C.** Extended haplotype homozygosity (EHH) plot around the lactoferrin for the R47 (blue) and K47 (red) around the variable position site, showing the extended haplotype around the K47 variant. **D.** Haplotype bifurcation plot showing breakdown of linkage disequilibrium in individuals carrying the lactoferrin R47 (blue) and K47 (red) alleles around the variant position. Thickness of the line corresponds to the number of individuals with shared haplotypes.

Given our observed correlations between sites under selection in primates and variation within human populations, we sought to determine whether lactoferrin exhibits signatures of positive selection among modern humans. Calculation of pairwise F_ST_ between a subset of human populations identified an elevated signal of differentiation between European (CEU) and African (YRI) populations [29], consistent with observed differences in allele frequencies between these groups (Figure S2A). The F_ST_ at rs1126478 was 0.70 (empirical p-value < 0.001), 0.30, and 0.03 for CEU-YRI, CEU-CHB, and CEU-FIN, respectively. Single nucleotide variants neighboring rs1126478 also showed signs of elevated F_ST_ suggesting that a shared CEU haplotype was driving the signal of differentiation (Figure S2A).

We next applied measures of haplotype homozygosity to assess the possibility that the K47 haplotype has been subject to natural selection in humans. Linkage around R47 alleles breaks down within ~10 kilobases, while the K47 variant possesses a markedly extended haplotype, consistent with the possibility of an adaptive sweep in this genomic region (Figure 2C). A selective sweep is also consistent with bifurcation plots around position 47, where the K47 haplotypes possess increased homogeneity relative to R47 haplotypes (Figure 2D). We observed a slight an elevation of the genome-wide corrected integrated haplotype score (iHS) for the K47 allele (-1.40136) and a depletion of observed heterozygosity (Figure S2B, S3, S4). We also examined the patterns of cross population extended haplotype homozygosity (XP-EHH). Consistent with the FST and EHH results, the XP-EHH score was elevated at the K47 position when CEU individuals were compared against YRI (1.1; p-value: 0.129) or CHB (3.1; p-value:>0.003)(Figure S5). While XP-EHH between CEU and YRI was moderate, surrounding SNPs less than 3 kilobases away had values as high as 2.89 (rs189460549; p-value: 0.01). Genome-wide, the K47 XP-EHH signal is moderate compared to other loci. Next we compared the joint distribution of the p-values from *d*N/*d*S analyses [24] with the empirical p-values from the CEU-CHB XP-EHH analyses (Figure S6). The previous genome-wide rank for lactoferrin, from *d*N/*d*S analyses, was 226 before considering the joint distribution and 156 after. The top 20 genes with the greatest change in rank (*d*N/*d*S p-value < 0.01) include *BLK*, *DSG1*, *FAS*, *SLC15A1*, *GLMN*, *SULT1C3*, *WIPF1*, *and LTF.* This meta-analysis highlights genes that have undergone species-level as well as population-level selection in primates and humans respectively. By integrating molecular phylogenetic analyses and population genetics approaches, we pinpointed signatures of positive selection associated with an abundant human lactoferrin polymorphism.

### Rapid evolution of lactoferrin-derived antimicrobial peptides

Evidence of positive selection in the lactoferrin N lobe among diverse primates, including position 47 in humans, led us to more closely investigate evolutionary pressures that have influenced variation in this region. After gene duplication from ancestral transferrin, lactoferrin gained potent antimicrobial activities independent of iron binding through cationic domains capable of disrupting microbial membranes. Two portions of the lactoferrin N lobe in particular, termed lactoferricin (amino acids 20-67) and lactoferrampin (amino acids 288-304), have been implicated in these antimicrobial functions [18,30]. Recent reports indicate that the human lactoferrin K47 variant, within the N lobe lactoferricin peptide, may have a protective effect against dental cavities associated with pathogenic bacteria [31]. Moreover, saliva isolated with patients homozygous for the K47 variant possesses enhanced antibacterial activity against oral *Streptococci* relative to homozygous R47 individuals [32].

Phylogenetic analysis revealed that several sites corresponding to lactoferricin and lactoferrampin display signatures of positive selection (Figure 3A, 3B). Notably, positive selection in lactoferricin localized to sites harboring cationic (lysine, arginine) or polar uncharged residues (asparagine), which could mediate membrane disruption and regulate antimicrobial activity. Position 47, which exhibits signatures of selection in humans as well as other primates, also lies within the lactoferricin peptide region. In contrast, hydrophobic tryptophan residues proposed to mediate insertion into microbial membranes are completely conserved among primates, as are cysteine residues that participate in intramolecular disulfide bond formation (Figure 3A). We also observed rapid evolution of the position immediately C-terminal to the cleavage site in lactoferrampin (Figure 3A), suggesting that the precise cleavage site in this peptide may be variable among species. Expanding our phylogenetic analysis to other mammalian taxa, we found that lactoferrin also exhibits signatures of positive selection in rodents and carnivores (Figure S7,Supplementary Table 10). While the specific positions that contribute most strongly to these signatures could not be resolved with high confidence, N-terminal regions corresponding to lactoferricin in primates are absent in several rodent and carnivore transcripts, suggesting that this activity may have been lost or modified in divergent mammals. Together these results demonstrate that lactoferrin-derived cationic peptides of the N lobe are rapidly evolving at sites critical for antimicrobial action.

**Figure 3.**
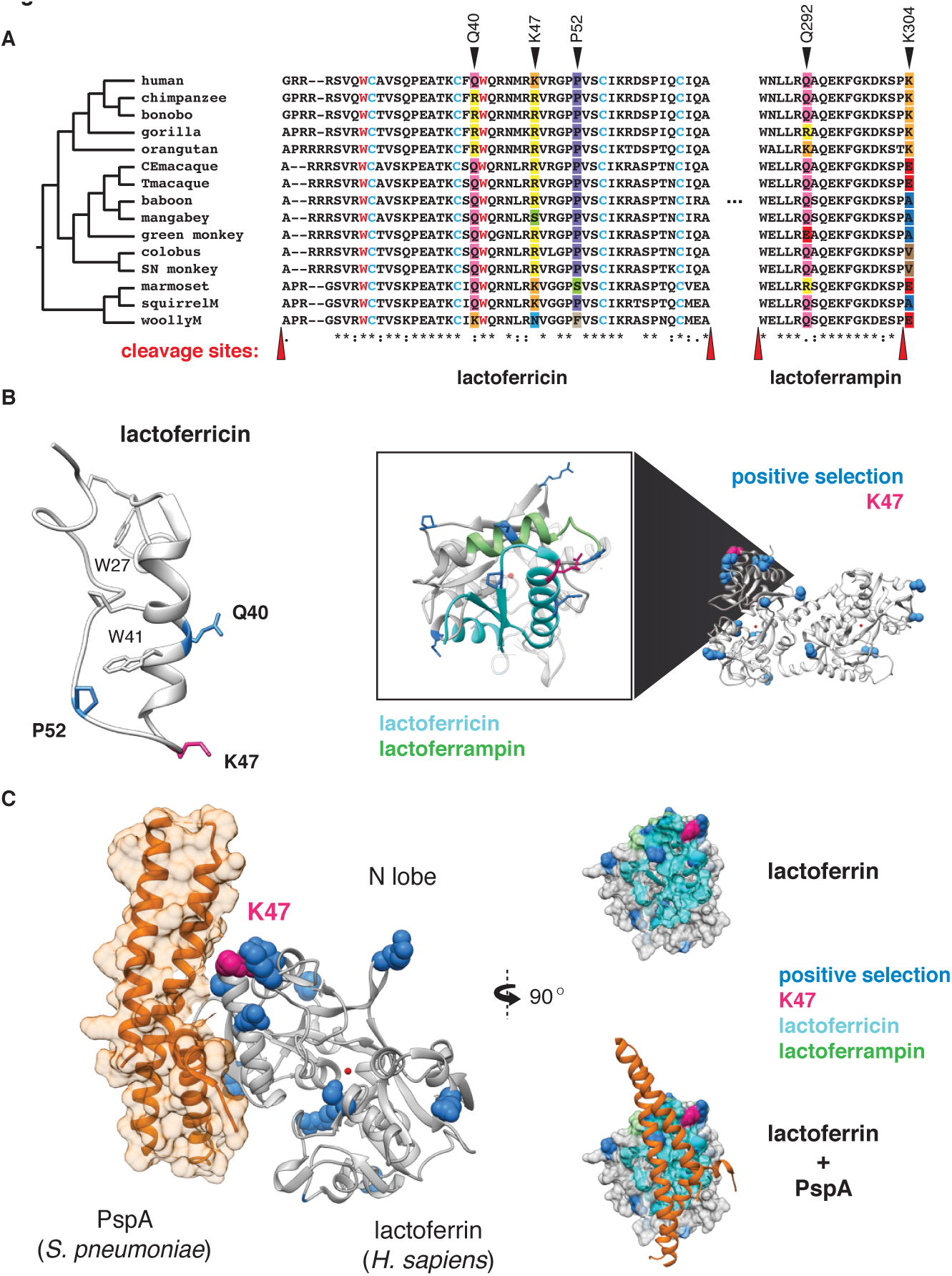
Rapid evolution of lactoferrin-derived antimicrobial peptides and pathogen binding interfaces. **A.** Amino acid alignment of the lactoferricin and lactoferrampin peptide sequences across primates. Sites under positive selection are denoted with black arrows, with amino acids at these positions color-coded. Conserved tryptophan (red) and cysteine (blue) residues are highlighted, which contribute to target membrane interactions and disulfide bond formation respectively. The reported cleavage sites of the two peptides are denoted with red arrows. **B.** Left: solution structure of the free human lactoferricin peptide (PDB: 1Z6V), with sites under positive selection (blue), including position 47 (magenta) indicated. Conserved tryptophan and cysteine residues highlighted in **A** are also shown. Right: enlarged view of the human lactoferrin N lobe highlighting sequences corresponding to lactoferricin (cyan) and lactoferrampin (green) antimicrobial peptides. Sites previously identified under positive selection in primates are shown in blue, with the position 47 variant shown in magenta. **C.** Crystal structure (PDB: 2PMS) of human lactoferrin N lobe (gray) bound to PspA from *Streptococcus pneumoniae* (orange). Side chains of sites under positive selection (blue), including position 47 (magenta) are shown.

## Distinct microbial interfaces are subject to positive selection in lactoferrin

While rapid evolution of the lactoferrin N lobe may reflect selection for improved targeting of pathogen surfaces, it could also represent adaptations that prevent binding by inhibitors encoded by pathogens. For example, pneumococcal surface protein A (PspA) is a crucial virulence determinant of *Streptococcus pneumoniae*, and several studies have demonstrated that PspA specifically binds and inhibits antimicrobial portions of the lactoferrin N lobe [33]. Consistent with an important evolutionary impact for this interaction, numerous sites under positive selection in the lactoferrin N lobe lie proximal to the PspA binding interface [34], including those corresponding to the lactoferricin peptide (Figure 3C). These data suggest that adaptive substitutions in lactoferrin could negate PspA binding, leading to enhanced immunity against *S. pneumoniae* or related pathogens.

Many strains of pathogenic *Neisseria*, which cause the sexually transmitted disease gonorrhea as well as acute meningitis, encode lactoferrin binding proteins (LbpA and LbpB) which mediate iron acquisition from lactoferrin [35,36]. Of four sites identified under positive selection in the lactoferrin C lobe, at least two appear proximal to the proposed *Neisseria* LbpA binding interface based on recent molecular modeling studies (Figure S8) [37]. One of these, position 589, also aligns to a region under strong positive selection in transferrin (position 576 in humans) which directly contacts the related bacterial receptor TbpA (Figure 1B) [28]. These findings suggest that, similarly to transferrin, antagonism by bacterial Lbp proteins may have promoted natural selection in the lactoferrin C lobe. Signatures of selection at distinct lactoferrin-pathogen interfaces thus highlight the diverse conflicts that have arisen during the evolution of this unique immunity factor.

## DISCUSSION

Together our results suggest that the emergence of novel antimicrobial activity in the N lobe of lactoferrin strongly influenced host-pathogen interactions in primates, including modern humans (Figure 4). High disparity in sites under positive selection between the N and C lobes of lactoferrin and transferrin indicate that distinct selective pressures influenced these proteins during primate evolution. We previously demonstrated that primate transferrin has been engaged in recurrent evolutionary conflicts with the bacterial receptor, TbpA [25]. This receptor is an important virulence factor in several Gram-negative human pathogens including *Neisseria gonorrhoeae*, *Neisseria meningitidis*, *Haemophilus influenzae*, as well as related animal pathogens [26,38–40]. Notably, TbpA binds and extracts iron exclusively from the C lobe of transferrin, and signatures of positive selection in transferrin are almost entirely restricted to the TbpA binding interface (Figure 1) [25]. The fact that transferrin family proteins are recurrently targeted by pathogens for iron acquisition may have provided the selective advantage for antimicrobial functions that arose in the lactoferrin N lobe.

**Figure 4.**
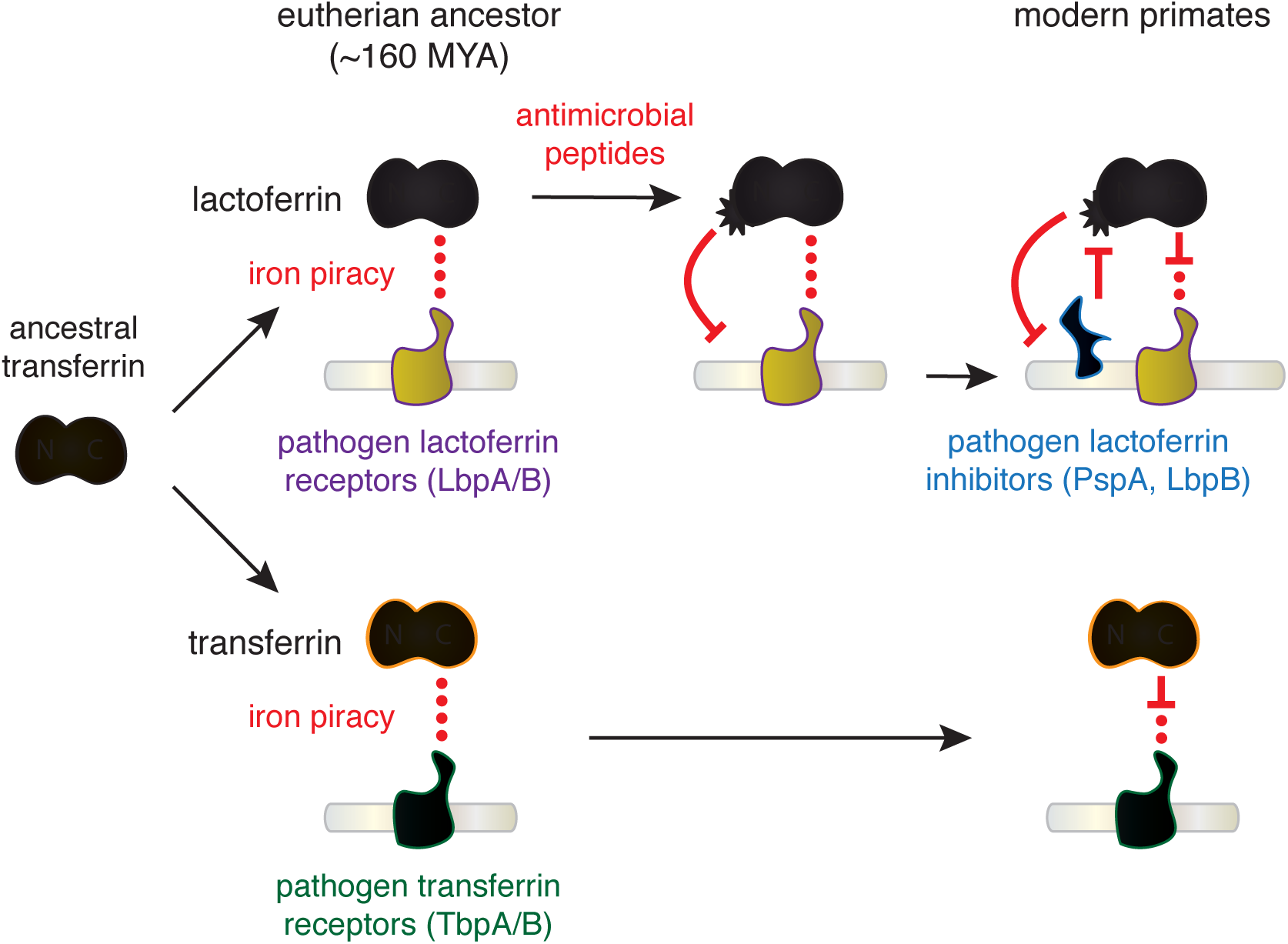
Model of lactoferrin evolution and genetic conflicts with pathogens. Following a duplication of the transferrin gene in the ancestor of eutherian mammals, interactions between the transferrin (yellow) C lobe and the bacterial transferrin receptors such as TbpA (green) led to the emergence of a molecular arms race.In contrast, while lactoferrin has likely also been engaged in evolutionary conflicts with pathogen iron acquisition receptors like LbpA (purple), the emergence of antimicrobial peptide activity in the N lobe would have provided novel defense activity against pathogens targeting lactoferrin as an iron source. This function would have led to the emergence of pathogen inhibitors of lactoferrin antimicrobial peptide activity (such as PspA or LbpB), which have dominated subsequent evolutionary conflicts localized to the lactoferrin N lobe.

Our results suggest at least two non-mutually exclusive scenarios for evolutionary conflicts involving the lactoferrin N lobe. Positive selection in this region could reflect adaption of lactoferrin for enhanced targeting of variable pathogen surfaces. Lactoferricin is capable of binding the bacterial LPS, which itself is heavily modified in many pathogens to mediate immune evasion and could provoke counter-adaptations at this interface. Conversely, variation in the lactoferrin N lobe could negate interactions with pathogen inhibitory proteins such as PspA encoded by *S. pneumoniae.* Lactoferrin binding activity has also been identified in several other important bacterial pathogens including *Treponema pallidum* [41], *Staphlococcus aureus* [42], and *Shigella flexneri* [43], raising the possibility of multiple independent evolutionary conflicts playing out at the lactoferrin N lobe. Iron-loaded lactoferrin could further be viewed as a “Trojan horse,” where microbes that target it as a nutrient iron source may be more susceptible to antimicrobial peptides. Consistent with this hypothesis, recent work has suggested that *Neisseria* encoded LbpB recognizes the lactoferrin N lobe, in contrast to its homolog TbpB which selectively interacts with the iron-loaded C lobe of transferrin [36,44,45]. LbpB binding to the lactoferrin N lobe could thus provide a counter-adaptation with dual benefits by neutralizing lactoferrin antimicrobial activity through negatively charged protein surfaces while simultaneously promoting iron acquisition by its co-receptor, LbpA [44]. These observations point to host-pathogen adaptations involving *de novo* protein functions on both sides of this genetic conflict.

Our results raise the possibility that the lactoferrin K47 variant introgressed into humans from Neanderthals at some point after the out-of-Africa expansion roughly 60,000 years ago [46]. An alternative explanation could be convergent evolution of lactoferrin in distinct lineages of early hominins for enhanced immune function, as well as long term balancing selection.Future analysis of lactoferrin sequence in archaic humans could provide additional insight on the history of this variant. Together these studies provide a direct link between variation in the lactoferrin N lobe and protection against pathogenic bacteria, consistent with adaptive evolution of lactoferrin in humans and other primates.

Notably, the lactoferrin gene, *LTF*, is located only ~60 kb away from *CCR5*, a chemokine receptor which is also an entry receptor for HIV [47–51]. A 32-base pair deletion in *CCR5* (*CCR5-Δ32*) confers resistance to HIV infection, and is present at a high frequency in northern Europeans while absent from African populations [52]. Although early evidence suggested that *CCR5-Δ32* might itself be subject to positive selection in humans, more recent studies have concluded that these signatures are more consistent with neutral evolution [53]. It is intriguing that, like *CCR5-Δ32*, the lactoferrin K47 variant exhibits increased allele frequency in European populations relative to Africans. However, the presence of the K47 variant at high frequencies in Asian and American populations points to a much earlier origin for this variant than *CCR5-Δ32*. Moreover, EHH and bifurcation analyses indicate that the haplotypes associated with the lactoferrin K47 variant do not encompass *CCR5*, suggesting that variation at the *CCR5* locus is unlikely to contribute to signatures of selection in *LTF* (Figure 2B, 2C, Supplementary Table 9). The proximity of the *LTF* and *CCR5* genes combined with their high degree of polymorphism and shared roles in immunity suggest the potential for genetic interactions relating to host defense. Future studies could reveal functional or epidemiological links between these two factors in human immunity.

In summary, we have discovered that lactoferrin constitutes a crucial node of host-pathogen evolutionary conflict based on signatures of natural selection across primates, including modern humans. Our findings suggest an intriguing mechanism for molecular arms race dynamics where adaptations and counter-adaptations rapidly emerge at the level of new protein functions in addition to recurrent amino acid substitutions at a single protein interface (Figure 4). Duplication and neofunctionalization of gene paralogs carries the potential to drastically alter the outcome of genetic conflicts. Our evolutionary analyses highlighting consequences of lactoferrin diversity pinpoint critical genetic variants to pursue in the fight against modern infectious diseases.

## MATERIALS & METHODS

### Primate genetic sources

RNA was obtained from the following species via the Coriell Cell Repositories where sample codes are indicated: *Homo sapiens* (human; primary human foreskin fibroblasts; gift from A. Geballe), *Gorilla gorilla* (western lowland gorilla; AG05251), *Papio anubis* (olive baboon; PR00036), *Lophocebus albigena* (grey-cheeked mangabey; PR01215),*Cercopithecus aethiops* (African green monkey; PR01193), *Colobus guereza* (colobus monkey; PR00240), *Callithrix geoffroyi* (white-fronted marmoset; PR00789), *Lagothrix lagotricha* (common woolly monkey; AG05356), *Saimiri sciureus* (common squirrel monkey; AG05311). Gene sequences from additional primate, rodent, and carnivore species were obtained from Genbank.

### cDNA cloning and sequencing

RNA (50 ng) from each primate cell line was prepared (RNeasy kit; Qiagen) and used as template for RT–PCR (SuperScript III; Invitrogen). Primers used to amplify lactoferrin cDNA were as follows: GTGGCAGAGCCTTCGTTTGCC (LF-forward; oMFB256) and GACAGCAGGGAATTGTGAGCAGATG (LF-rev; oMFB313). PCR products were TA-cloned into pCR2.1 (Invitrogen) and directly sequenced from at least three individual clones. Gene sequences have been deposited in Genbank (KT006751 – KT006756).

### Phylogenetic analyses and structural observations

DNA multiple sequence alignments were performed using MUSCLE and indels were manually trimmed based on amino-acid comparisons. A generally accepted primate species phylogeny [54] (Figure 1A) was used for evolutionary analysis. A gene tree generated from the alignment of lactoferrin corresponded to this species phylogeny (PhyML; http://atgc.lirmm.fr/phyml/). Maximum-likelihood analysis of the lactoferrin and transferrin data sets was performed with codeml of the PAML software package [21]. A free-ratio model allowing *d*N/*d*S (omega) variation along branches of the phylogeny was employed to calculate *d*N/*d*S values between lineages. Two-ratio tests were performed using likelihood models to compare all branches fixed at *d*N/*d*S=1 or an average *d*N/*d*S value from the whole tree applied to each branch to varying *d*N/*d*S values according to branch.

Positive selection in lactoferrin was assessed by fitting the multiple alignment to either F3X4 or F61 codon frequency models. Likelihood ratio tests (LRTs) were performed by comparing pairs of site-specific models (NS sites): M1 (neutral) with M2 (selection), M7 (neutral, beta distribution of *d*N/*d*S<1) with M8 (selection, beta distribution, *d*N/*d*S>1 allowed). Additional LRTs from the HyPhy software package that also account for synonymous rate variation and recombination (FUBAR, REL, FEL, MEME, BUSTED) were performed [22,23]. Molecular structures of lactoferrin, transferrin and associated proteins were visualized using Chimera (http://www.cgl.ucsf.edu/chimera/).

### Human population genetics analysis

For variant-based analyses we used genotype calls from the 1000 Genomes project (release: 20130502, shapeit2 phased). Weir and Cockerham’s F_s_t estimator [29] was used for the population comparisons, implemented in GPAT++. EHH and the bifurcation diagrams were calculated using the [R] package REHH [55]. Genome-wide iHS1 scans were performed using GPAT++ and XPEHH plots were generated previously published datasets [56,57].

## ACKNOWLEDGEMENTS

The authors thank members of the Elde lab and N. Barber for helpful discussions and comments on the manuscript. This work was supported by the National Institutes of Health awards R01GM114514 (N.C.E.) and K99GM115822. N.C.E. is a Pew Scholar in the Biomedical Sciences and Mario R. Capecchi Endowed Chair in Genetics.

---

**Figure S1.**
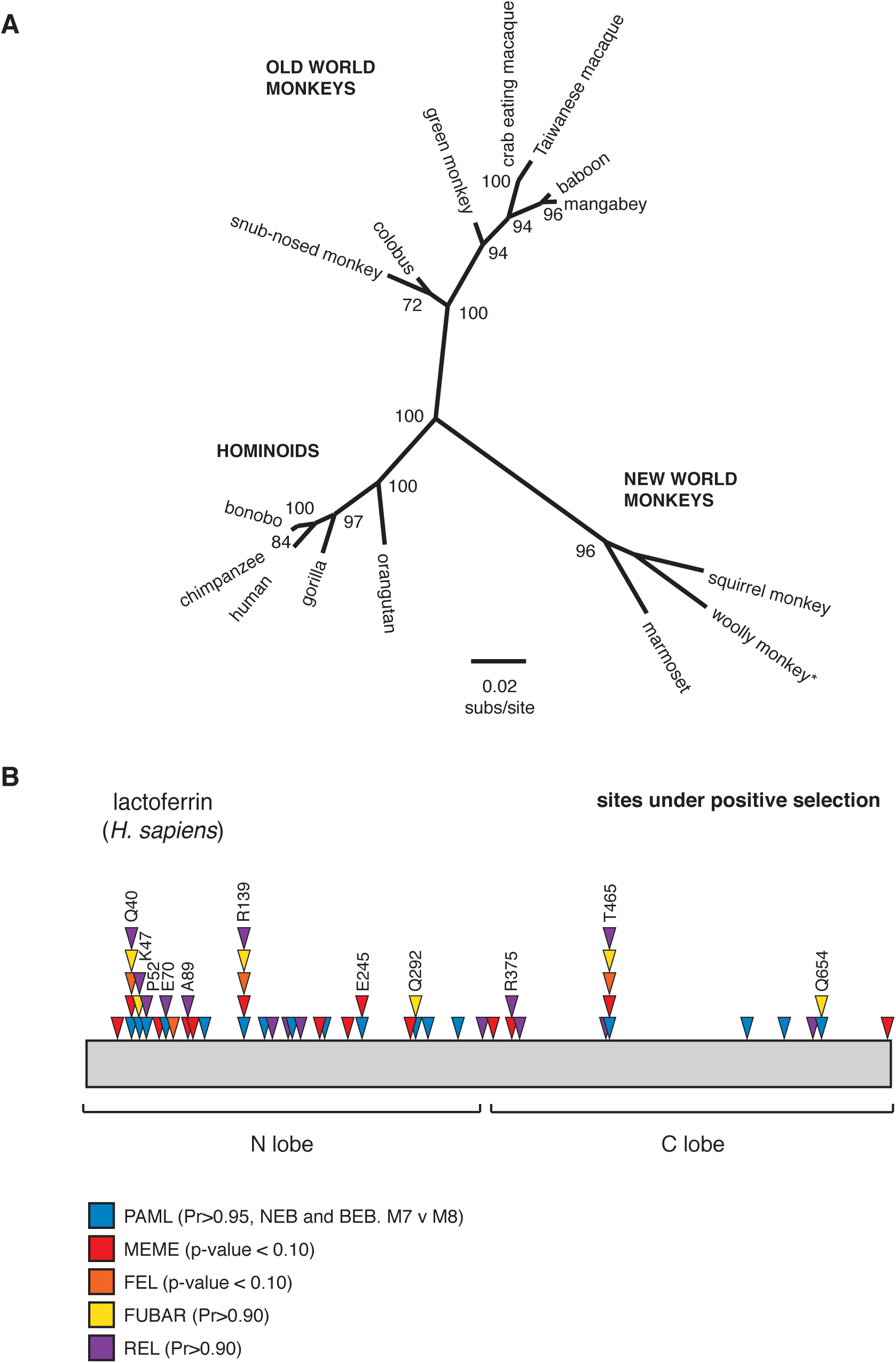
**A.** Phylogeny for primate lactoferrin gene (*LTF*) inferred using PhyML with the LG substitution model. Bootstrap values are indicated at branch nodes. Astericks denotes orientation of New World monkeys, which differs slightly from the accepted species phylogeny (Figure 1A). **B.** Sites identified under positive selection using PAML and HyPhy analyses. Positions identified by at least two separate tests for selection are indicated.

**Figure S2.**
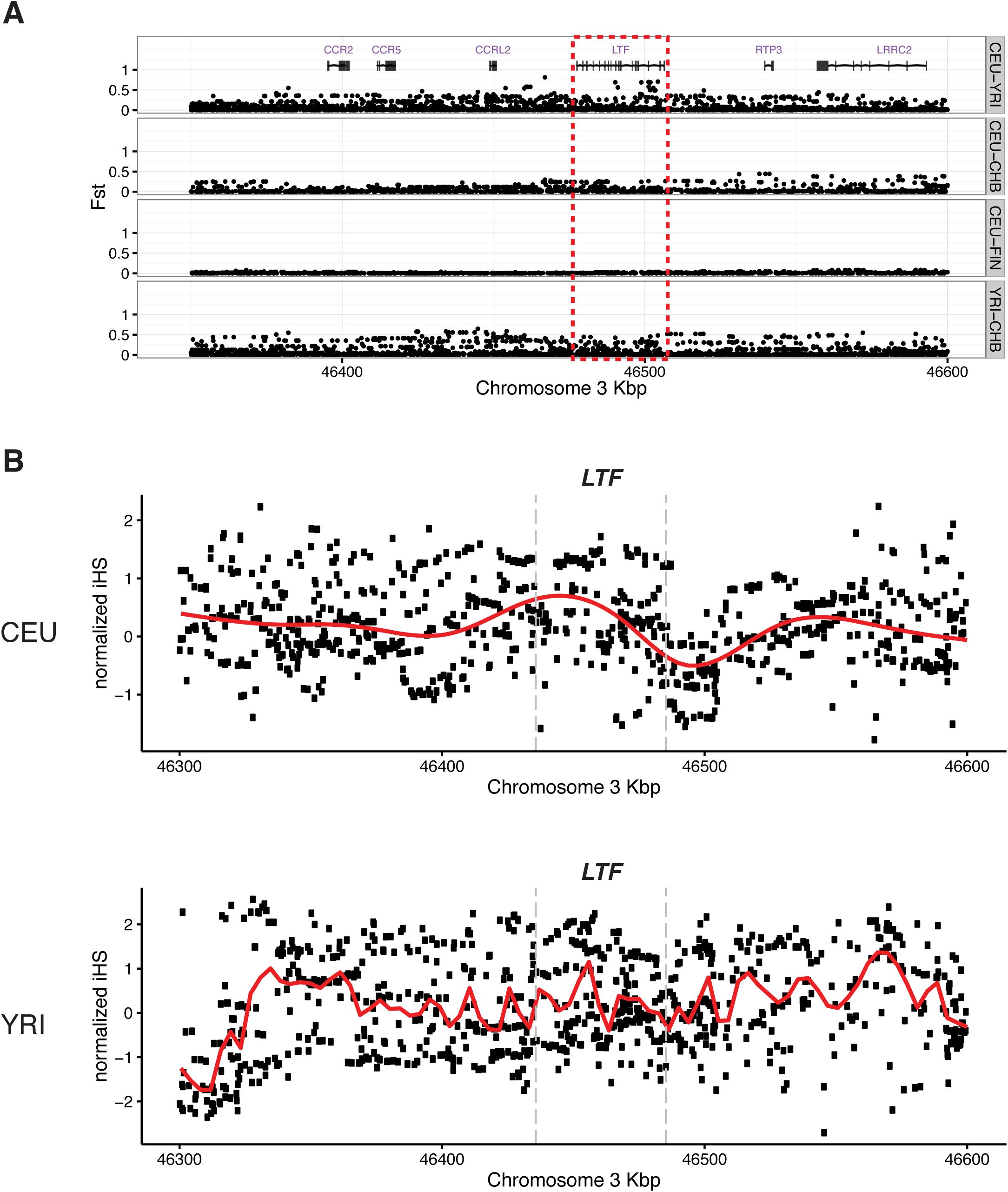
**A.** Pairwise Fst calculated between indicated human populations (CEU: Western European; YRI: Yoruban; CHB: Chinese; FIN: Finnish) proximal to the lactoferrin gene locus (*LTF).* The boundaries of the *LTF*gene are marked in red. **B.** Normalized iHS calculations at the *LTF*locus within CEU and YRI populations. Gray bars denote the boundaries of *LTF.* Red line represents the locally weighted scatterplot smoothing (LOESS) regression.

**Figure S3.**
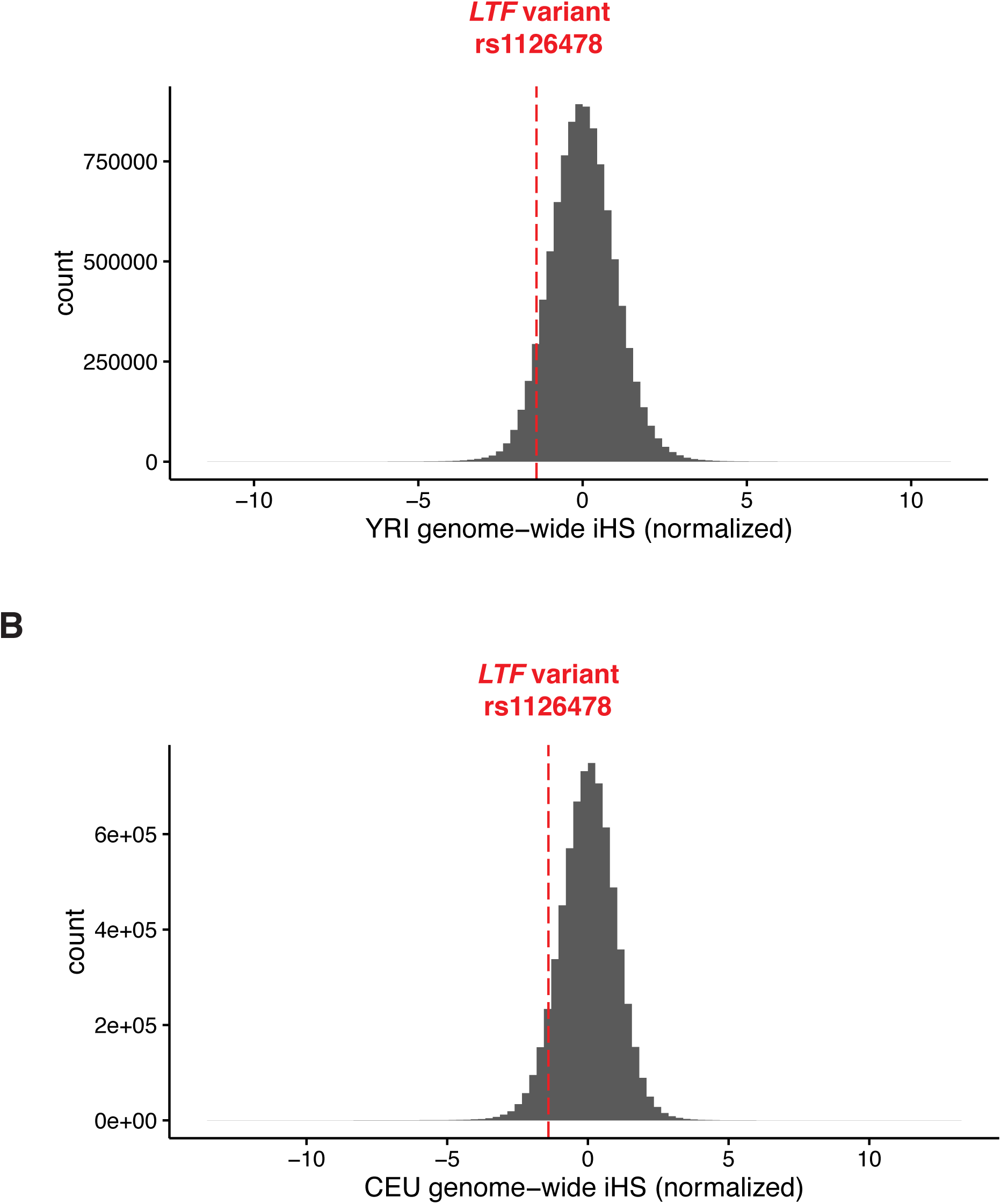
Histograms representing genome-wide iHS values within YRI (**A**) and CEU (**B**) populations calculated from One Thousand Genomes Project Phase III data. Negative values denote an extended haplotype in reference relative to non-reference alleles. Red line highlights the *LTF* amino acid position 47 variant (rs1126478).

**Figure S4.**
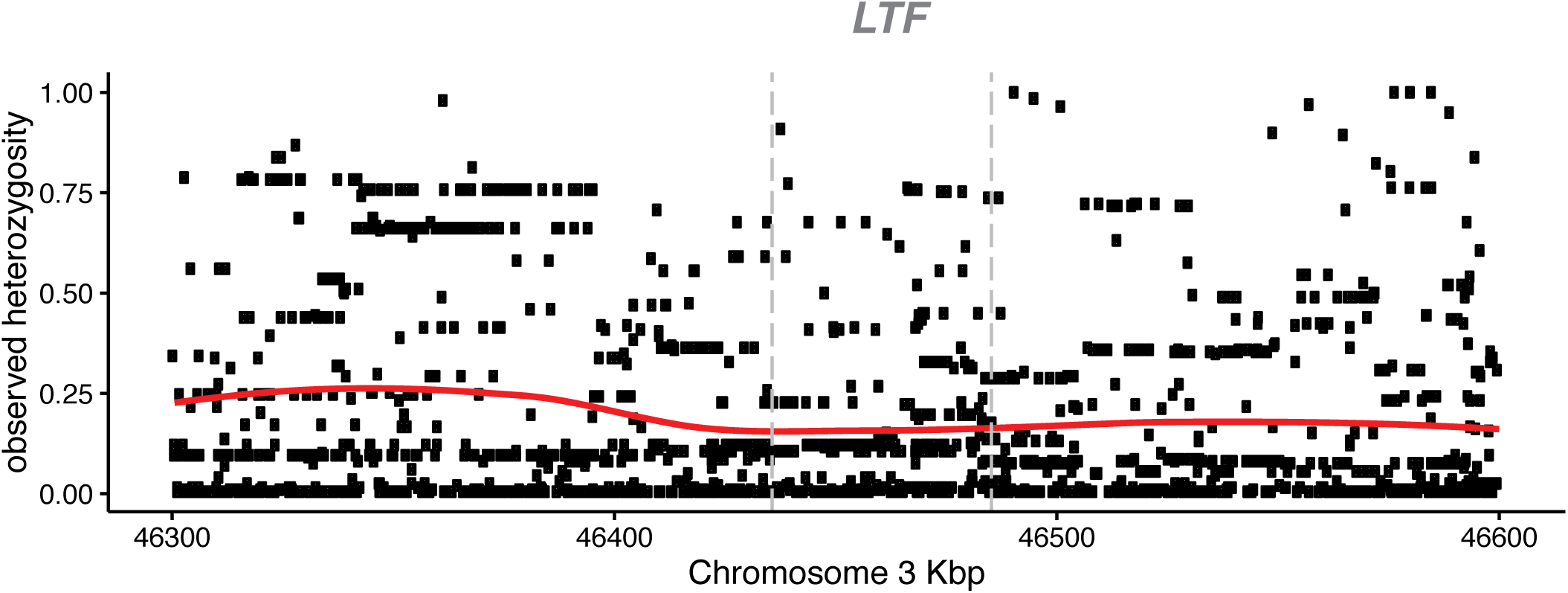
Observed heterozygosity at the *LTF* locus. Observed heterozygosity is measured as the fraction of heterozygous genotypes at a variant position. Gray lines denote the start and end of *LTF.*

**Figure S5.**
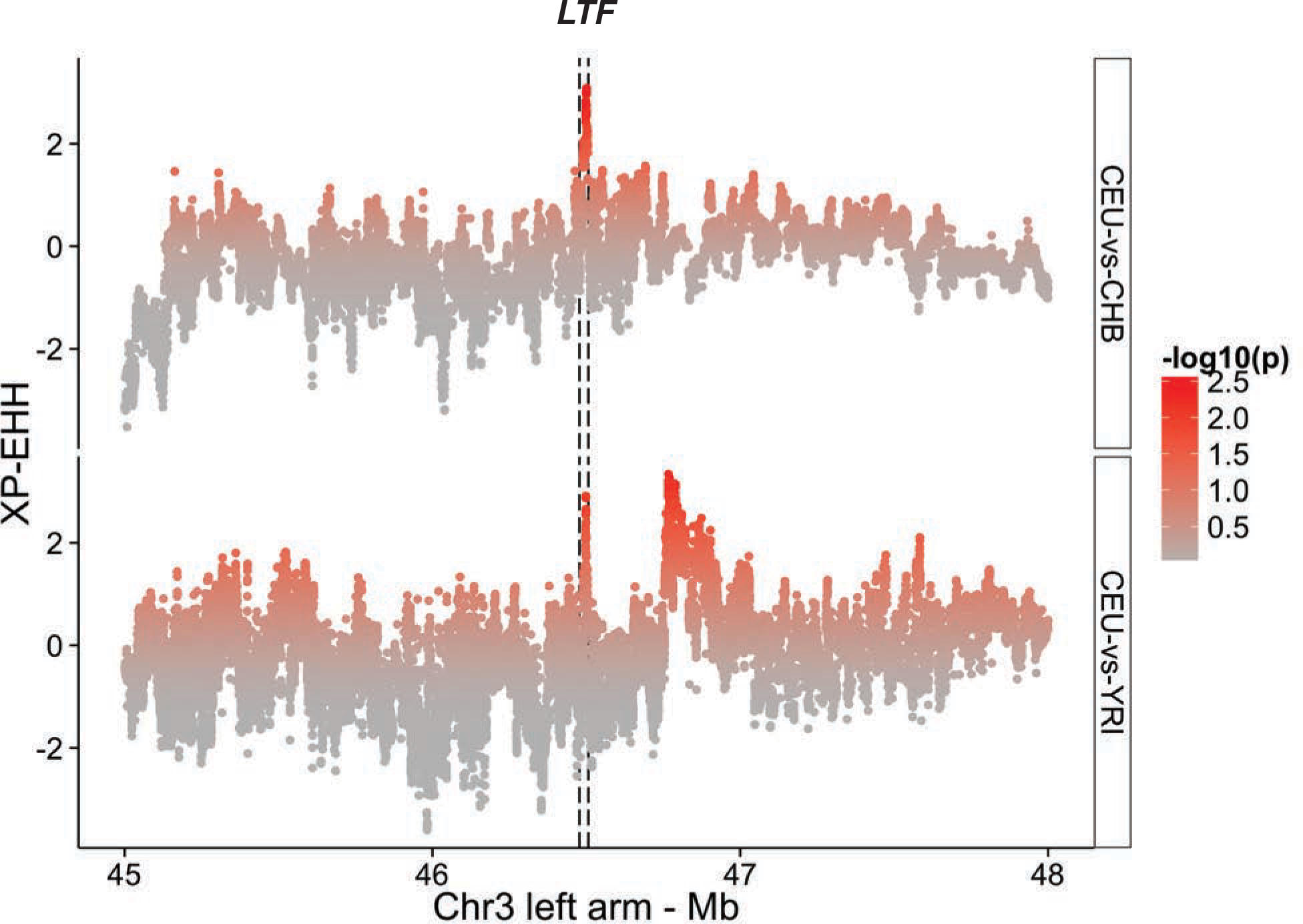
Elevated XP-EHH signal between human populations (CEU and YRI or CHB) at the *LTF* locus. Dashed lines highlights the boundaries of the *LTF* gene. P-values were empirically determined by Prybus et al. 2014.

**Figure S6.**
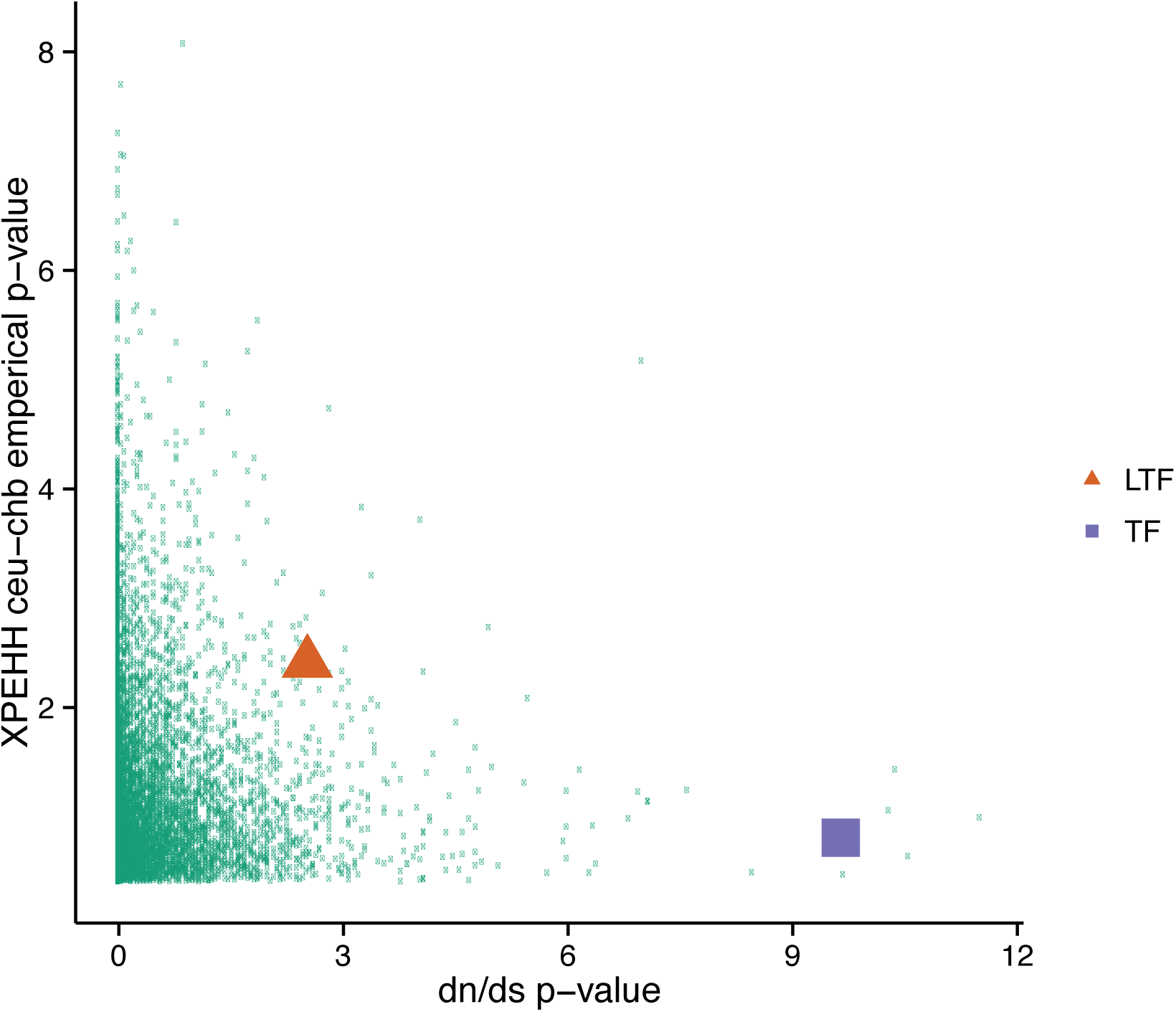
The joint distribution of the *d*N/*d*S p-value (George et al., 2011) and the empirical CEU-CHB p-value (Prybus et al. 2014). Single genes are denoted by green dots. Lactoferrin (LTF) and transferrin (TF) are highlighted.

**Figure S7.**
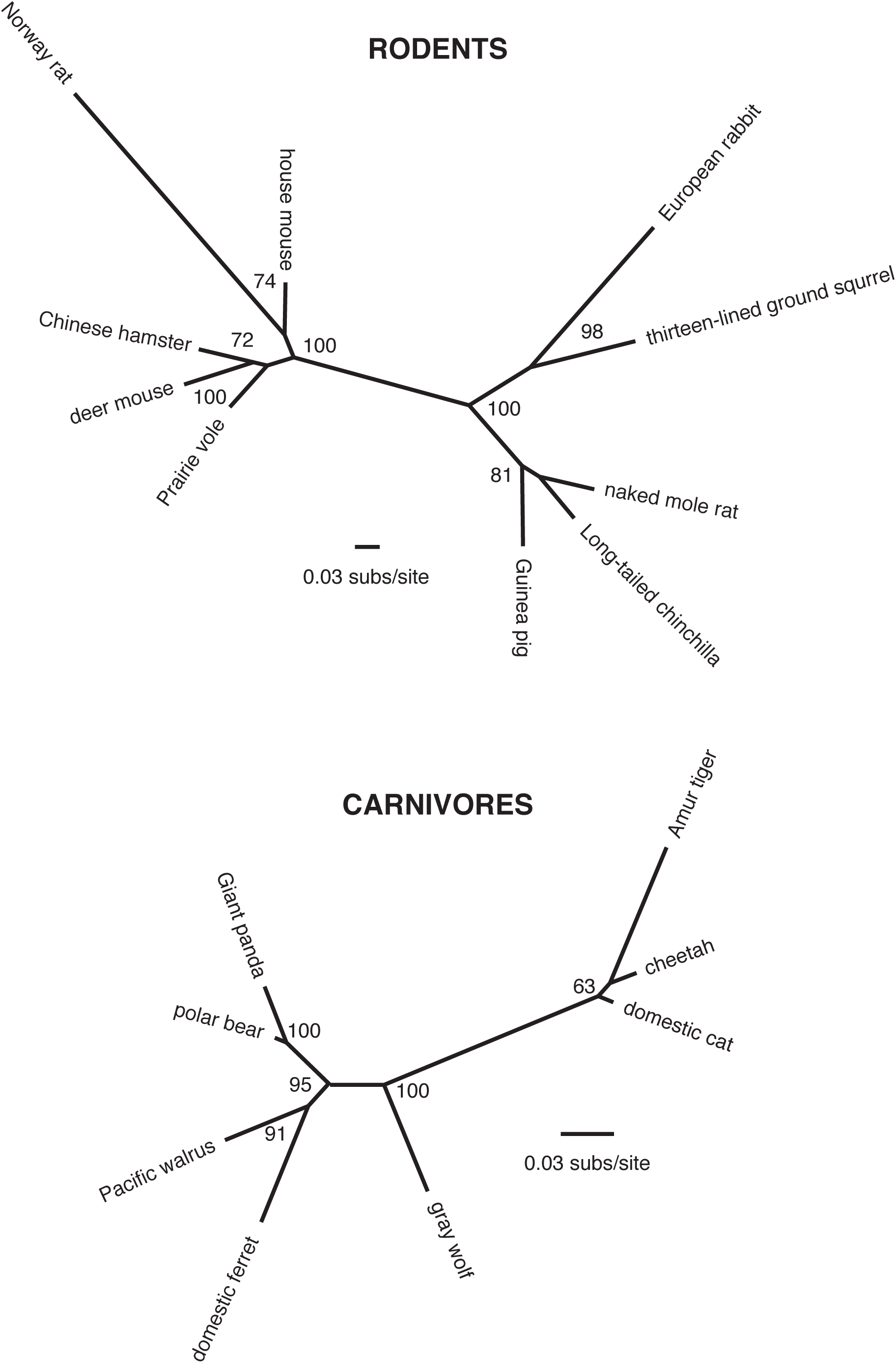
Phylogenies for rodent and carnivore LTF genes inferred using PhyML. Bootstrap values are indicated at branch nodes.

**Figure S8.**
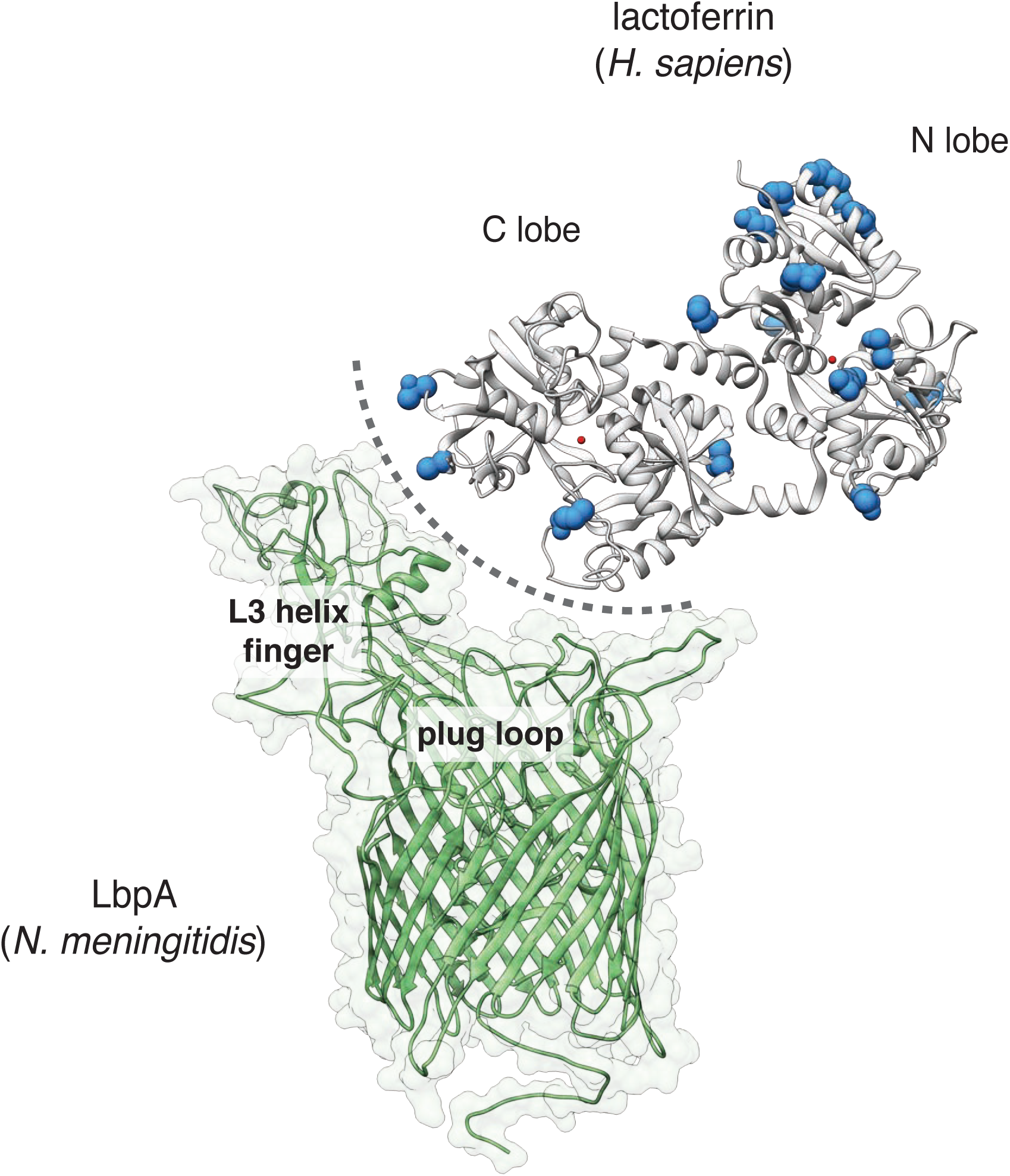
Signatures of positive selection at the proposed lactoferrin-LbpA binding interface (Noinaj et al., 2013). Crystal structure for lactoferrin is oriented to a model of the *N. meningitidis* LbpA protein inferred from structure of the related protein TbpA. Sites subject to positive selection in primate lactoferrin are marked in blue. The proposed binding interface is denoted with a dashed line. Positions of the LbpA plug loop and L3 helix finger, which contribute to iron acquistion in the homologous protein TbpA, are marked.

**Supplementary Table 1.** Likelihood ratio test statistics for models of variable selection along branches of the primate lactoferrin phylogeny using PAML.

|  | **lnL\*** | **2δ** | **Df\*\*** | **p-value** |
| --- | --- | --- | --- | --- |
| Model 0<br>Same dN/dS for all branches | -6357.95 |  |  |  |
| Model 1<br>Different dN/dS ratio for different branches | -6267.33 | 181.24 | 26 | <0.0001 |

\* likelihood score
\*\* degrees of freedom, equal to one less than the total number of branches in the phylogeny.

**Supplementary Table 2.** Summary of positive selection in primate lactoferrin using PAML. Analyses were performed using two independent codon models (F3X4, F61) which describe the frequency and rate of substitutions. Selection was inferred by comparing likelihood scores between models that allow for selection (M2, M8) relative models which exclude selection (M1, M7) in this gene.

|  | Codon freq. | *M1-M2* |  | *M7-M8* |  | Tree length | dN/dS (%) |
| --- | --- | --- | --- | --- | --- | --- | --- |
| | | 2δ | p-value | 2δ | p-value | | |
| **Whole gene** | F3X4 | 16.9 | 0.00021 | 18.6 | <0.0001 | 0.90 | 1.8 (22.5) |
|  | F61 | 21.5 | <0.0001 | 23.0 | <0.0001 | 0.89 | 1.9 (22.2) |
| **N-lobe** | F3X4 | 14.3 | 0.00079 | 14.8 | 0.00060 | 0.99 | 2.1 (20.6) |
|  | F61 | 17.8 | 0.00014 | 18.1 | 0.00012 | 0.96 | 2.0 (26.7) |
| **C-lobe** | F3X4 | 4.2 | 0.12 | 4.3 | 0.12 | 0.81 | 1.6 (20.3) |
|  | F61 | 6.5 | 0.038 | 7.3 | 0.025 | 0.80 | 1.9 (17.2) |

**Supplementary Table 3.**
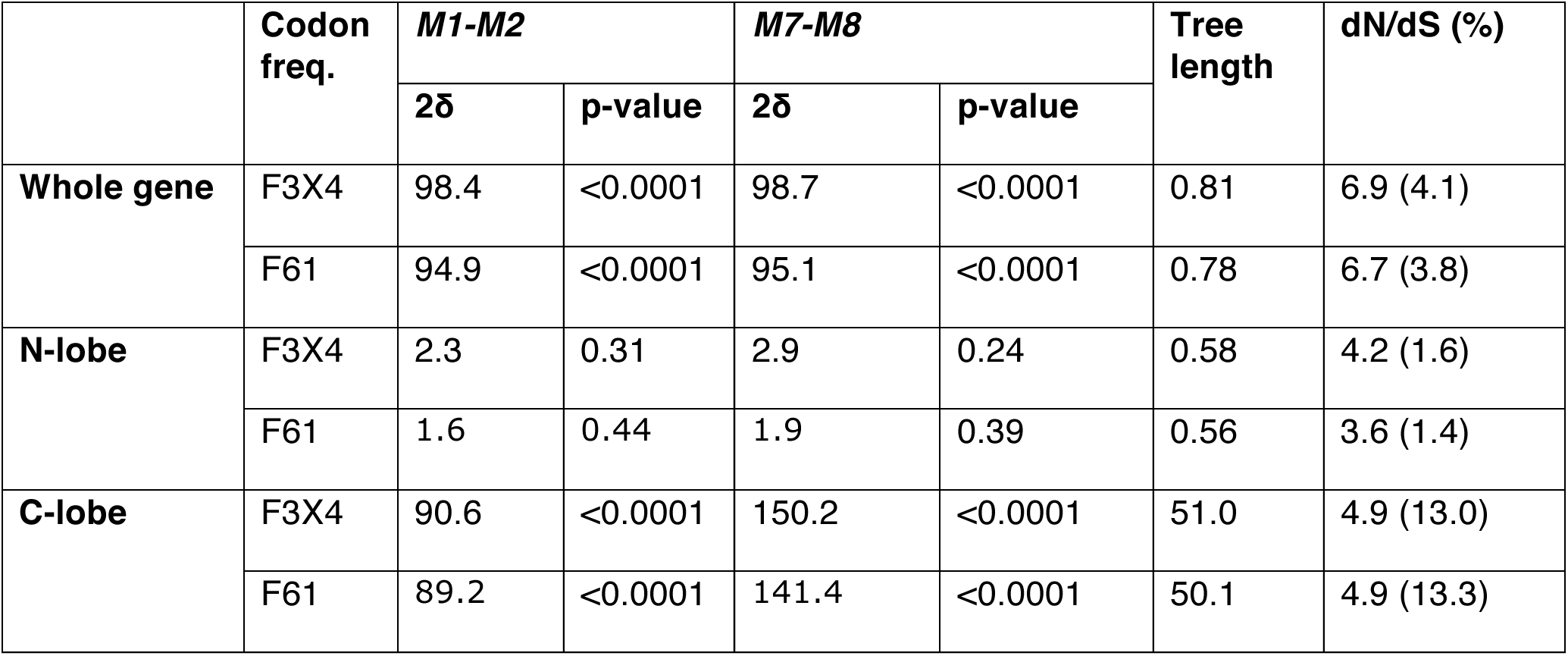
Summary of positive selection in primate transferrin using PAML. Analyses were performed using two independent codon models (F3X4, F61) which describe the frequency and rate of substitutions. Selection was inferred by comparing likelihood scores between models that allow for selection (M2, M8) relative models which exclude selection (M1, M7) in this gene.

|  | Codon freq. | *M1-M2* |  | *M7-M8* |  | Tree length | dN/dS (%) |
| --- | --- | --- | --- | --- | --- | --- | --- |
| | | 2δ | p-value | 2δ | p-value | | |
| **Whole gene** | F3X4 | 98.4 | <0.0001 | 98.7 | <0.0001 | 0.81 | 6.9 (4.1) |
|  | F61 | 94.9 | <0.0001 | 95.1 | <0.0001 | 0.78 | 6.7 (3.8) |
| **N-lobe** | F3X4 | 2.3 | 0.31 | 2.9 | 0.24 | 0.58 | 4.2 (1.6) |
|  | F61 | 1.6 | 0.44 | 1.9 | 0.39 | 0.56 | 3.6 (1.4) |
| **C-lobe** | F3X4 | 90.6 | <0.0001 | 150.2 | <0.0001 | 51.0 | 4.9 (13.0) |
|  | F61 | 89.2 | <0.0001 | 141.4 | <0.0001 | 50.1 | 4.9 (13.3) |

**Supplementary Table 4.** Lactoferrin whole gene log likelihood scores and parameter estimates for four models of variable ω among sites assuming the F3×4 model of codon frequencies (PAML). Amino acid positions shown are for human lactoferrin.

| Site model | Parameter estimates | Sites* with $\omega^{**}$ AA ☐ | lnL |
| --- | --- | --- | --- |
| **M1: neutral** | ($\omega_0=0$) $f_0=0.635$<br>($\omega_1=1$) $f_1=0.365$<br>branch $\omega$ (mean)= 0.365 | | -6267.33 |
| **M2: selection** | ($\omega_0=0.06$) $f_0=0.731$<br>($\omega_1=1$) $f_1=0.122$<br>**($\omega_2=2.1$) $f_2=0.146$**<br>branch $\omega$ (mean)= 0.466 | M623 | -6258.87 |
| **M7: $\beta$** | $p=0.005$<br>$q=0.0074$<br>branch $\omega$ (mean)= 0.40 | | -6268.21 |
| **M8: $\beta$ and $\omega$** | $p=8.415$<br>$q=99.00$ ($f_0=0.775$)<br>**$\omega_1=1.8$ ($f_1=0.225$)**<br>branch $\omega$ (mean)= 0.465 | Q40<br>K47<br>P52<br>E70<br>R105<br>R139<br>T158<br>G183<br>S210<br>E245<br>Q292<br>K304<br>P331<br>T465<br>A588<br>M622<br>Q654 | -6258.92 |

\* posterior probabilities >0.95 by both Naïve Empirical Bayes (NEB) and Bayes Empirical Bayes analysis.
\*\* $\omega = dN/dS$

**Supplementary Table 5.** Summary of positive selection in primate lactoferrin (MEME, FEL). Amino acid positions shown are for human lactoferrin.

| Model | Sites with evidence of positive selection (p-value) |
| --- | --- |
| MEME | Q26 (0.018)<br>Q40 (0.089)<br>Q63 (0.087)<br>A89 (0.097)<br>K92 (0.020)<br>R139 (0.00020)<br>E206 (0.0030)<br>E230 (0.084)<br>E245 (0.0087)<br>N288 (0.019)<br>R361 (0.050)<br>R375 (0.015)<br>T465 (0.035)<br>R709 (0.014) |
| FEL | Q40 (0.075)<br>V76 (0.097)<br>R139 (0.044)<br>T465 (0.034) |

**Supplementary Table 6.** Summary of positive selection in primate lactoferrin using FUBAR and REL algorithms. Amino acid positions shown are for human lactoferrin.

| Model | Sites with evidence of diversifying selection | Posterior probability | Bayes Factor |
| --- | --- | --- | --- |
| FUBAR | Q40 | 0.90 | 17.1 |
|  | K47 | 0.91 | 18.3 |
|  | R139 | 0.96 | 49.2 |
|  | Q292 | 0.93 | 23.6 |
|  | T465 | 0.97 | 64.6 |
|  | Q654 | 0.92 | 22.5 |
| REL | Q40 | 0.94 | 67.6 |
|  | K47 | 0.93 | 57.7 |
|  | P52 | 0.94 | 72.1 |
|  | E70 | 0.95 | 83.0 |
|  | A89 | 0.93 | 61.4 |
|  | R139 | 0.94 | 71.2 |
|  | P163 | 0.94 | 67.9 |
|  | K182 | 0.93 | 57.6 |
|  | G183 | 0.95 | 77.3 |
|  | A193 | 0.92 | 51.9 |
|  | K352 | 0.92 | 52.1 |
|  | R375 | 0.93 | 60.0 |
|  | G382 | 0.93 | 58.6 |
|  | R462 | 0.92 | 52.2 |
|  | T465 | 0.95 | 77.4 |
|  | D648 | 0.93 | 61.6 |

**Supplementary Table 7.** BUSTED likelihood ratio test statistics for gene-wide episodic diversifying selection in primate lactoferrin.

| Model | log L | AICc | Tree length | LRT p-value |
| --- | --- | --- | --- | --- |
| Unconstrained | -6305.60 | 12703.61 | 0.91 | 0.001 |
| Constrained | -6312.57 | 12715.52 | 0.86 |  |

**Supplementary Table 8.**
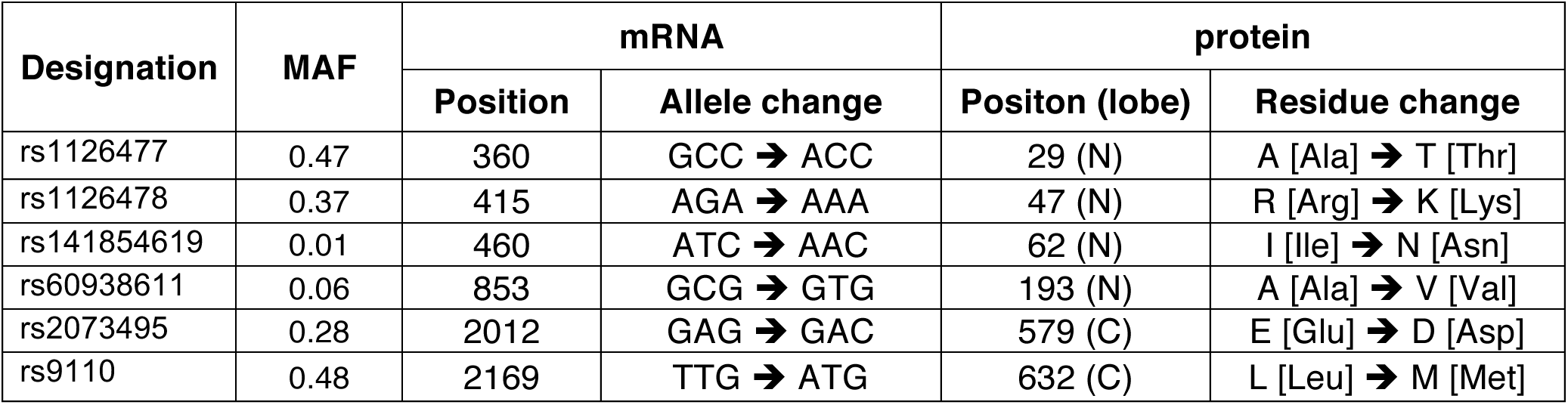
Summary of abundant lactoferrin missense variants in human population. Allele frequencies are reported from 1000 Genomes project data (phase 3). MAF: minor allele frequency.

| Designation | MAF | mRNA |  | protein |  |
| --- | --- | --- | --- | --- | --- |
|  |  | Position | Allele change | Positon (lobe) | Residue change |
| rs1126477 | 0.47 | 360 | GCC → ACC | 29 (N) | A [Ala] → T [Thr] |
| rs1126478 | 0.37 | 415 | AGA → AAA | 47 (N) | R [Arg] → K [Lys] |
| rs141854619 | 0.01 | 460 | ATC → AAC | 62 (N) | I [Ile] → N [Asn] |
| rs60938611 | 0.06 | 853 | GCG → GTG | 193 (N) | A [Ala] → V [Val] |
| rs2073495 | 0.28 | 2012 | GAG → GAC | 579 (C) | E [Glu] → D [Asp] |
| rs9110 | 0.48 | 2169 | TTG → ATG | 632 (C) | L [Leu] → M [Met] |

**Supplementary Table 9.**
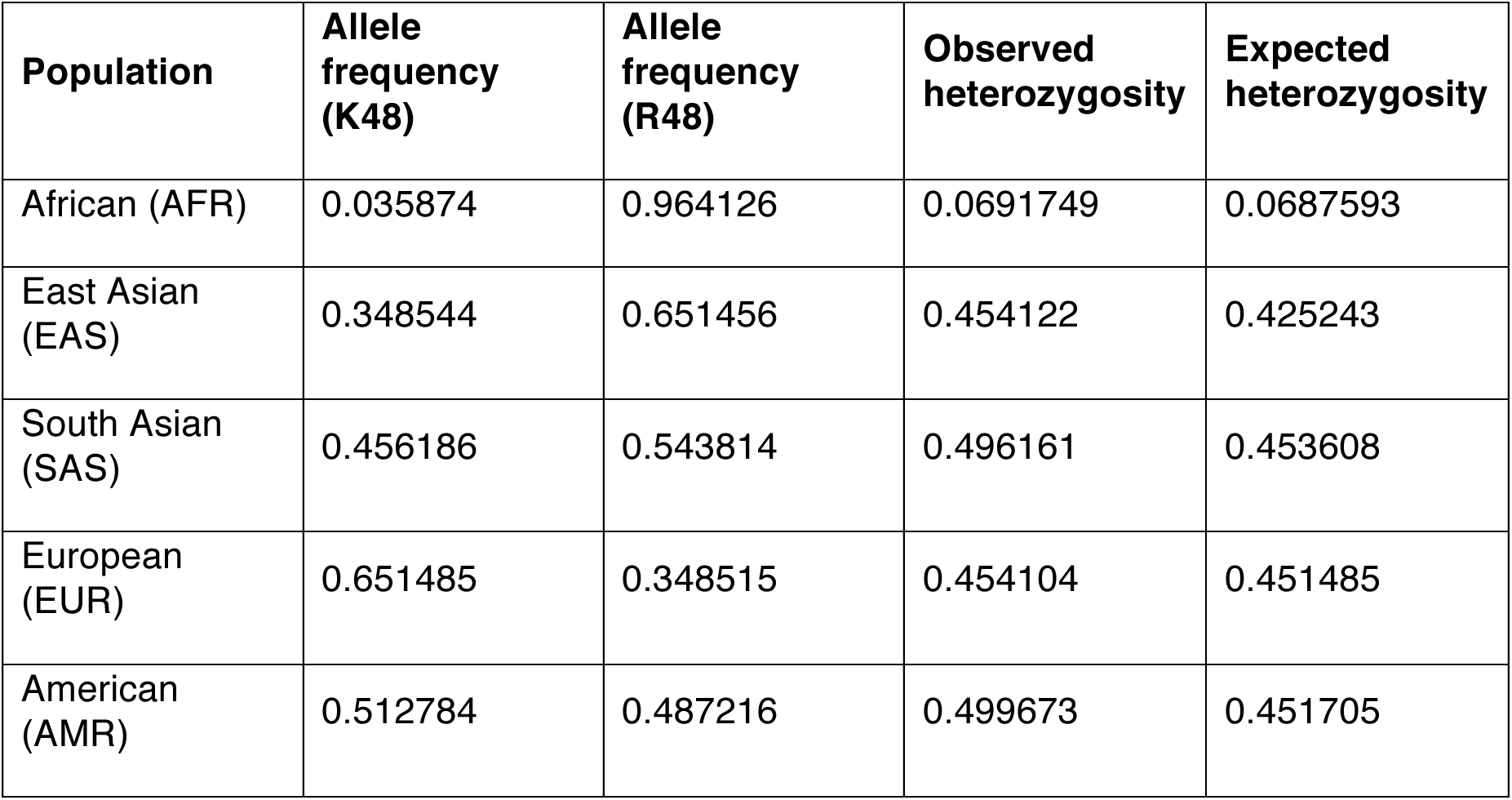
Allele frequencies for lactoferrin rs1126478 variant in 1000 Genomes database (phase 3).

| Population | Allele frequency (K48) | Allele frequency (R48) | Observed heterozygosity | Expected heterozygosity |
| --- | --- | --- | --- | --- |
| African (AFR) | 0.035874 | 0.964126 | 0.0691749 | 0.0687593 |
| East Asian (EAS) | 0.348544 | 0.651456 | 0.454122 | 0.425243 |
| South Asian (SAS) | 0.456186 | 0.543814 | 0.496161 | 0.453608 |
| European (EUR) | 0.651485 | 0.348515 | 0.454104 | 0.451485 |
| American (AMR) | 0.512784 | 0.487216 | 0.499673 | 0.451705 |

**Supplementary Table 10.** Summary of positive selection in rodent and carnivore lactoferrin using PAML. Analyses were performed using two independent codon models (F3×4, F61). Selection was inferred by comparing likelihood scores between models that allow for selection (M2, M8) relative models which exclude selection (M1, M7) in this gene. Tree length and dN/dS values are shown for the M8 calculations.

|  | Codon freq. | *M1-M2* |  | *M7-M8* |  | Tree length | dN/dS (%) |
| --- | --- | --- | --- | --- | --- | --- | --- |
| | | 2δ | p-value | 2δ | p-value | | |
| **Rodents**<br>**(10 species)** | F3X4 | 9.0 | 0.011 | 31.0 | <0.0001 | 4.1 | 1.9 (5.1) |
|  | F61 | 0 | 1.0 | 24.3 | <0.0001 | 3.9 | 2.1 (4.1) |
| **Carnivores**<br>**(8 species)** | F3X4 | 6.6 | 0.037 | 9.7 | 0.0079 | 1.1 | 4.3 (1.7) |
|  | F61 | 4.5 | 0.10 | 5.7 | 0.059 | 1.1 | 4.5 (1.2) |

## REFERENCES

1. Daugherty MD, Malik HS. Rules of Engagement: Molecular Insights from Host-Virus Arms Races. Annu Rev Genet. 2012;46: 677–700. doi:10.1146/annurev-genet-110711-155522

2. Haldane J. Disease and evolution. La Ricerca Scientifica Supplemento. 1949;: 1–11.

3. Van Valen L. A new evolutionary law. Evol Theory. 1973;1: 1–30.

4. Hamilton WD, Axelrod R, Tanese R. Sexual reproduction as an adaptation to resist parasites (a review). Proc Natl Acad Sci USA. 1st ed. National Academy of Sciences; 1990;87: 3566–3573.

5. Sawyer SL, Wu LI, Emerman M, Malik HS. Positive selection of primate TRIM5α identifies a critical species-specific retroviral restriction domain. Proc Natl Acad Sci USA. National Acad Sciences; 2005;102: 2832–2837. doi:10.1073/pnas.0409853102

6. Elde NC, Child SJ, Geballe AP, Malik HS. Protein kinase R reveals an evolutionary model for defeating viral mimicry. Nature. 2009;457: 485–489. doi:10.1038/nature07529

7. Tenthorey JL, Kofoed EM, Daugherty MD, Malik HS, Vance RE. Molecular basis for specific recognition of bacterial ligands by NAIP/NLRC4 inflammasomes. Molecular Cell. 2014;54: 17–29. doi:10.1016/j.molcel.2014.02.018

8. Barber MF, Elde NC. Buried Treasure: Evolutionary Perspectives on Microbial Iron Piracy. Trends in Genetics. Elsevier Ltd; 2015;31: 627–636. doi:10.1016/j.tig.2015.09.001

9. Lambert LA. Molecular evolution of the transferrin family and associated receptors. Biochimica et Biophysica Acta. 2012;1820: 244–255. doi:10.1016/j.bbagen.2011.06.002

10. Weinberg ED. Nutritional immunity. Hosts attempt to withold iron from microbial invaders. JAMA. 1975;231: 39–41.

11. Hood MI, Skaar EP. Nutritional immunity: transition metals at the pathogen-host interface. Nat Rev Micro. 2012;10: 525–537. doi:10.1038/nrmicro2836

12. Cassat JE, Skaar EP. Iron in infection and immunity. 2013;13: 509–519. doi:10.1016/j.chom.2013.04.010

13. Morgenthau A, Pogoutse A, Adamiak P, Moraes TF, Schryvers AB. Bacterial receptors for host transferrin and lactoferrin: molecular mechanisms and role in host-microbe interactions. Future Microbiol. 2013;8: 1575–1585. doi:10.2217/fmb.13.125

14. Andrews NC. Disorders of iron metabolism. N Engl J Med. 1999;341: 1986–1995. doi:10.1056/NEJM199912233412607

15. Weinberg ED. Iron withholding: a defense against infection and neoplasia. Physiol Rev. 1984;64: 65–102.

16. García-Montoya IA, Cendón TS, Arévalo-Gallegos S, Rascón-Cruz Q. Lactoferrin a multiple bioactive protein: an overview. Biochimica et Biophysica Acta. 2012;1820: 226–236. doi:10.1016/j.bbagen.2011.06.018

17. Berlutti F, Pantanella F, Natalizi T, Frioni A, Paesano R, Polimeni A, et al. Antiviral properties of lactoferrin‐‐a natural immunity molecule. Molecules. 2011;16: 6992–7018. doi:10.3390/molecules16086992

18. Haney EF, Nazmi K, Lau F, Bolscher JGM, Vogel HJ. Novel lactoferrampin antimicrobial peptides derived from human lactoferrin. Biochimie. 2009;91: 141–154. doi:10.1016/j.biochi.2008.04.013

19. Elass-Rochard E, Roseanu A, Legrand D, Trif M, Salmon V, Motas C, et al. Lactoferrin-lipopolysaccharide interaction: involvement of the 28-34 loop region of human lactoferrin in the high-affinity binding to Escherichia coli 055B5 lipopolysaccharide. Biochem J. 1995;312 ( Pt 3): 839–845.

20. Vorland LH, Ulvatne H, Rekdal O, Svendsen JS. Initial binding sites of antimicrobial peptides in Staphylococcus aureus and Escherichia coli. Scand J Infect Dis. 1999;31: 467–473.

21. Yang Z. PAML 4: phylogenetic analysis by maximum likelihood. Molecular Biology and Evolution. Oxford University Press; 2007;24: 1586–1591. doi:10.1093/molbev/msm088

22. Pond SLK, Frost SDW, Muse SV. HyPhy: hypothesis testing using phylogenies. Bioinformatics. 2005;21: 676–679. doi:10.1093/bioinformatics/bti079

23. Delport W, Poon AFY, Frost SDW, Kosakovsky Pond SL. Datamonkey 2010: a suite of phylogenetic analysis tools for evolutionary biology. Bioinformatics. 2010;26: 2455–2457. doi:10.1093/bioinformatics/btq429

24. George RD, McVicker G, Diederich R, Ng SB, MacKenzie AP, Swanson WJ, et al. Trans genomic capture and sequencing of primate exomes reveals new targets of positive selection. Genome Research. Cold Spring Harbor Lab; 2011;21: 1686–1694. doi:10.1101/gr.121327.111

25. Barber MF, Elde NC. Escape from bacterial iron piracy through rapid evolution of transferrin. Science. 2014;346: 1362–1366. doi:10.1126/science.1257259

26. Cornelissen CN, Biswas GD, Tsai J, Paruchuri DK, Thompson SA, Sparling PF. Gonococcal transferrin-binding protein 1 is required for transferrin utilization and is homologous to TonB-dependent outer membrane receptors. journal of bacteriology. 1992;174: 5788–5797.

27. Moraes TF, Yu R-H, Strynadka NCJ, Schryvers AB. Insights into the Bacterial Transferrin Receptor: The Structure of Transferrin-Binding Protein B from Actinobacillus pleuropneumoniae. Molecular Cell. 2009;35: 523–533. doi:10.1016/j.molcel.2009.06.029

28. Noinaj N, Easley NC, Oke M, Mizuno N, Gumbart J, Boura E, et al. Structural basis for iron piracy by pathogenic Neisseria. Nature. 2012;483: 53–58. doi:10.1038/nature10823

29. Weir BS, Cockerham CC. Estimating F-statistics for the analysis of population structure. Evolution. 1984;38: 1358. doi:10.2307/2408641

30. Yamauchi K, Tomita M, Giehl TJ, Ellison RT. Antibacterial activity of lactoferrin and a pepsin-derived lactoferrin peptide fragment. Infection and Immunity. 1993;61: 719–728.

31. Azevedo LF, Pecharki GD, Brancher JA, Cordeiro CA, Medeiros KGDS, Antunes AA, et al. Analysis of the association between lactotransferrin (LTF) gene polymorphism and dental caries. J Appl Oral Sci. 2010;18: 166–170.

32. Fine DH, Toruner GA, Velliyagounder K, Sampathkumar V, Godboley D, Furgang D. A lactotransferrin single nucleotide polymorphism demonstrates biological activity that can reduce susceptibility to caries. Infection and Immunity. 2013;81: 1596–1605. doi:10.1128/IAI.01063-12

33. Shaper M, Hollingshead SK, Benjamin WH, Briles DE. PspA Protects Streptococcus pneumoniae from Killing by Apolactoferrin, and Antibody to PspA Enhances Killing of Pneumococci by Apolactoferrin. Infection and Immunity. 2004;72: 5031–5040. doi:10.1128/IAI.72.9.5031-5040.2004

34. Senkovich O, Cook WJ, Mirza S, Hollingshead SK, Protasevich II, Briles DE, et al. Structure of a complex of human lactoferrin N-lobe with pneumococcal surface protein a provides insight into microbial defense mechanism. Journal of Molecular Biology. 2007;370: 701–713. doi:10.1016/j.jmb.2007.04.075

35. Schryvers AB, Morris LJ. Identification and characterization of the human lactoferrin-binding protein from Neisseria meningitidis. Infection and Immunity. 1988;56: 1144–1149.

36. Brooks CL, Arutyunova E, Lemieux MJ. The structure of lactoferrin-binding protein B from Neisseria meningitidis suggests roles in iron acquisition and neutralization of host defences. Acta Crystallogr F Struct Biol Commun. 2014;70: 1312–1317. doi:10.1107/S2053230X14019372

37. Noinaj N, Cornelissen CN, Buchanan SK. Structural insight into the lactoferrin receptors from pathogenic Neisseria. Journal of Structural Biology. 2013;184: 83–92. doi:10.1016/j.jsb.2013.02.009

38. Cornelissen CN, Kelley M, Hobbs MM, Anderson JE, Cannon JG, Cohen MS, et al. The transferrin receptor expressed by gonococcal strain FA1090 is required for the experimental infection of human male volunteers. Molecular Microbiology. 1998;27: 611–616.

39. Gray-Owen SD, Loosmore S, Schryvers AB. Identification and characterization of genes encoding the human transferrin-binding proteins from Haemophilus influenzae. Infection and Immunity. American Society for Microbiology (ASM); 1995;63: 1201–1210.

40. Silva LP, Yu R, Calmettes C, Yang X, Moraes TF, Schryvers AB, et al. Conserved interaction between transferrin and transferrin-binding proteins from porcine pathogens. Journal of Biological Chemistry. 2011;286: 21353–21360. doi:10.1074/jbc.M111.226449

41. Deka RK, Brautigam CA, Tomson FL, Lumpkins SB, Tomchick DR, Machius M, et al. Crystal structure of the Tp34 (TP0971) lipoprotein of treponema pallidum: implications of its metal-bound state and affinity for human lactoferrin. J Biol Chem. 2007;282: 5944–5958. doi:10.1074/jbc.M610215200

42. Naidu AS, Miedzobrodzki J, Musser JM, Rosdahl VT, Hedström SA, Forsgren A. Human lactoferrin binding in clinical isolates of Staphylococcus aureus. J Med Microbiol. 1991;34: 323–328.

43. Tigyi Z, Kishore AR, Maeland JA, Forsgren A, Naidu AS. Lactoferrin-binding proteins in Shigella flexneri. Infection and Immunity. Am Soc Microbiol; 1992;60: 2619–2626.

44. Morgenthau A, Beddek A, Schryvers AB. The negatively charged regions of lactoferrin binding protein B, an adaptation against anti-microbial peptides. PLOS one. 2014. doi:10.1371/journal.pone.0086243.t002

45. Yu RH, Schryvers AB. Regions located in both the N-lobe and C-lobe of human lactoferrin participate in the binding interaction with bacterial lactoferrin receptors. Microb Pathog. 1993;14: 343–353. doi:10.1006/mpat.1993.1034

46. Ségurel L, Quintana-Murci L. Preserving immune diversity through ancient inheritance and admixture. Current Opinion in Immunology. 2014;30: 79–84. doi:10.1016/j.coi.2014.08.002

47. Deng H, Liu R, Ellmeier W, Choe S, Unutmaz D, Burkhart M, et al. Identification of a major co-receptor for primary isolates of HIV-1. Nature. 1996;381: 661–666. doi:10.1038/381661a0

48. Dragic T, Litwin V, Allaway GP, Martin SR, Huang Y, Nagashima KA, et al. HIV-1 entry into CD4+ cells is mediated by the chemokine receptor CC-CKR-5. Nature. 1996;381: 667–673. doi:10.1038/381667a0

49. Doranz BJ, Rucker J, Yi Y, Smyth RJ, Samson M, Peiper SC, et al. A dual-tropic primary HIV-1 isolate that uses fusin and the beta-chemokine receptors CKR-5, CKR-3, and CKR-2b as fusion cofactors. 1996;85: 1149–1158.

50. Alkhatib G, Combadiere C, Broder CC, Feng Y, Kennedy PE, Murphy PM, et al. CC CKR5: a RANTES, MIP-1alpha, MIP-1beta receptor as a fusion cofactor for macrophage-tropic HIV-1. Science. 1996;272: 1955–1958.

51. Choe H, Farzan M, Sun Y, Sullivan N, Rollins B, Ponath PD, et al. The beta-chemokine receptors CCR3 and CCR5 facilitate infection by primary HIV-1 isolates. 1996;85: 1135–1148.

52. Samson M, Libert F, Doranz BJ, Rucker J, Liesnard C, Farber CM, et al. Resistance to HIV-1 infection in caucasian individuals bearing mutant alleles of the CCR-5 chemokine receptor gene. Nature. 1996;382: 722–725. doi:10.1038/382722a0

53. Sabeti PC, Walsh E, Schaffner SF, Varilly P, Fry B, Hutcheson HB, et al. The case for selection at CCR5-Δ32. Plos Biology. 2005;3. doi:10.1371/journal.pbio.0030378.st005

54. Perelman P, Johnson WE, Roos C, Seuánez HN, Horvath JE, Moreira MAM, et al. A molecular phylogeny of living primates. Plos Genetics. 2011;7: e1001342. doi:10.1371/journal.pgen.1001342

55. Gautier M, Vitalis R. rehh: an R package to detect footprints of selection in genome-wide SNP data from haplotype structure. Bioinformatics. 2012;28: 1176–1177. doi:10.1093/bioinformatics/bts115

56. Prendergast J, Maclean CA, Chue Hong N. hapbin: An efficient program for performing haplotype based scans for positive selection in large genomic datasets. Dataset. 2015;http://dx.doi.org/10.7488/ds/214.

57. Pybus M, Dall'Olio GM, Luisi P, Uzkudun M, Carreño-Torres A, Pavlidis P, et al. 1000 Genomes Selection Browser 1.0: a genome browser dedicated to signatures of natural selection in modern humans. Nucleic Acids Research. 2014;42: D903–9. doi:10.1093/nar/gkt1188

